# Honey bee queens are vulnerable to heat-induced loss of fertility

**DOI:** 10.1101/627729

**Authors:** Alison McAfee, Abigail Chapman, Heather Higo, Robyn Underwood, Joseph Milone, Leonard J Foster, M Marta Guarna, David R Tarpy, Jeffery S Pettis

## Abstract

All species need to reproduce to maintain viable populations, but heat stress kills sperm cells across the animal kingdom and rising frequencies of heatwaves are a threat to biodiversity. Honey bees (*Apis mellifera*) are globally distributed micro-livestock; therefore, they could serve as environmental biomonitors for fertility losses. Here, we found that queens have two potential routes of temperature-stress exposure: within colonies and during routine shipping. Our data suggest that temperatures of 15 to 38°C are safe for queens at a tolerance threshold of 11.5% loss of sperm viability, which is the viability difference associated with queen failure in the field. Heat shock activates expression of specific stress-response proteins in the spermatheca, which could serve as molecular biomarkers (indicators) for heat stress. This protein fingerprint may eventually enable surveys for the prevalence of heat-induced loss of sperm viability in diverse landscapes as part of a biomonitoring program.

---

Climate change is threatening biodiversity around the globe^1–3^, and one potential driver is through heat-induced reductions in fertility^4–6^. The impact of heat on fertility is far-reaching in the animal kingdom, affecting mammals^7–11^, birds^12^, fish^13^, nematodes^14^, and insects^4,15–21^. At temperatures of 40-43 LC, spermatogenesis is compromised^22^, sperm viability drops^4,18,19^, sperm are less competitive^4^, and motility is compromised^15,23,24^. Extreme weather events such as heatwaves are increasing in frequency and severity^25–27^, which could have widespread effects on populations via reduced reproductive output^4–6^.

Insects and other ectothermic animals are especially vulnerable to changes in local temperatures because, unlike mammals, they are less able to thermoregulate. Insects and are critical components of ecosystems and agriculture, with economic estimates placing the global value of insect pollination at approximately €153 billion annually^28^. Worryingly, declines in insect populations^29^ and pollinating species^30–34^ have already been reported, with habitat destruction, range compression, pesticide residues, diseases, and their interactions commonly reported as potential drivers^28,35,36^. However, there has been relatively little research on the effects of heat on pollinating insect reproduction^18,19^.

Young honey bee queens have one brief mating period early in life, and store the sperm they acquire for the rest of their lives (up to 5 years)^37^. As the sole egg-layer, colony productivity directly depends on the queen’s reproductive output, which in turn depends on the viability and abundance of her stored sperm^18,38,39^.

Like other insects, individual honey bees are ectothermic, but colonies can thermoregulate and maintain stable core temperatures of around 35°C in the brood nest^40^. Although colonies can persist in regions with extreme heat, evidence suggests that extreme ambient temperatures (38 - 46°C) are associated with colony losses^41^. Furthermore, internal hive temperatures positively correlate with ambient temperatures above 18°C, and brood nest temperatures can rise upwards of 37°C, both under natural conditions^42^ and during simulated heatwaves^43^. Hives occasionally experience spikes of > 40°C^44^, suggesting that queens could be vulnerable to temperature extremes even inside the hive.

Honey bees have excellent potential for being temperature stress biomonitors. Although we do not expect them to be sensitive sentinels, other attributes make honey bees amenable to a biomonitoring program: they are a globally distributed, managed species, so they are readily available in almost any geographic region, and they are already accepted as effective biomonitors for pollution^45^. If we begin to observe signs of heat stress in honey bees colonies, that would signal a worrisome risk of reduced reproductive output for ectothermic species.

Here, we begin to explore honey bees’ utility as temperature stress biomonitors. We monitored temperature fluctuations in colonies under extreme weather conditions, establishing that damaging intra-hive temperatures can occur. Next, we tested a range of temperatures and exposure durations to determine thresholds above which queen quality is likely to be compromised. We then investigated the biochemical basis of heat-induced sperm viability reduction in queens and drones using quantitative proteomics, which showed how heat stress alters protein expression of reproductive tissues. The specific set of upregulated proteins we identified may eventually serve as diagnostic tools to elucidate causes of queen failure and eventually enable regional surveys of heat stress as part of an environmental temperature biomonitoring program.

## Results and Discussion

### Sperm viability losses associated with queen failure

To establish how much of a reduction in sperm viability is associated with field-observable reduced reproductive output (and associated economic losses for beekeepers), we collected queens rated as ‘failing’ (N = 58) and ‘healthy’ (N = 55) by beekeepers and measured the queens’ stored sperm viability (**Fig 1a**). We found that the failed and healthy viability data were normally distributed (Shapiro test, P = 0.18 and 0.11, respectively), and that failed queens had significantly lower sperm viability (Student’s t test, P = 5.8E-06, F = 28.16), with an average drop of 11.5%. We then set an 11.5% viability drop as the tolerance threshold in subsequent experiments aimed at identifying critical temperatures beyond which queens are at risk of substantial loss of stored sperm viability.

**Figure 1.**
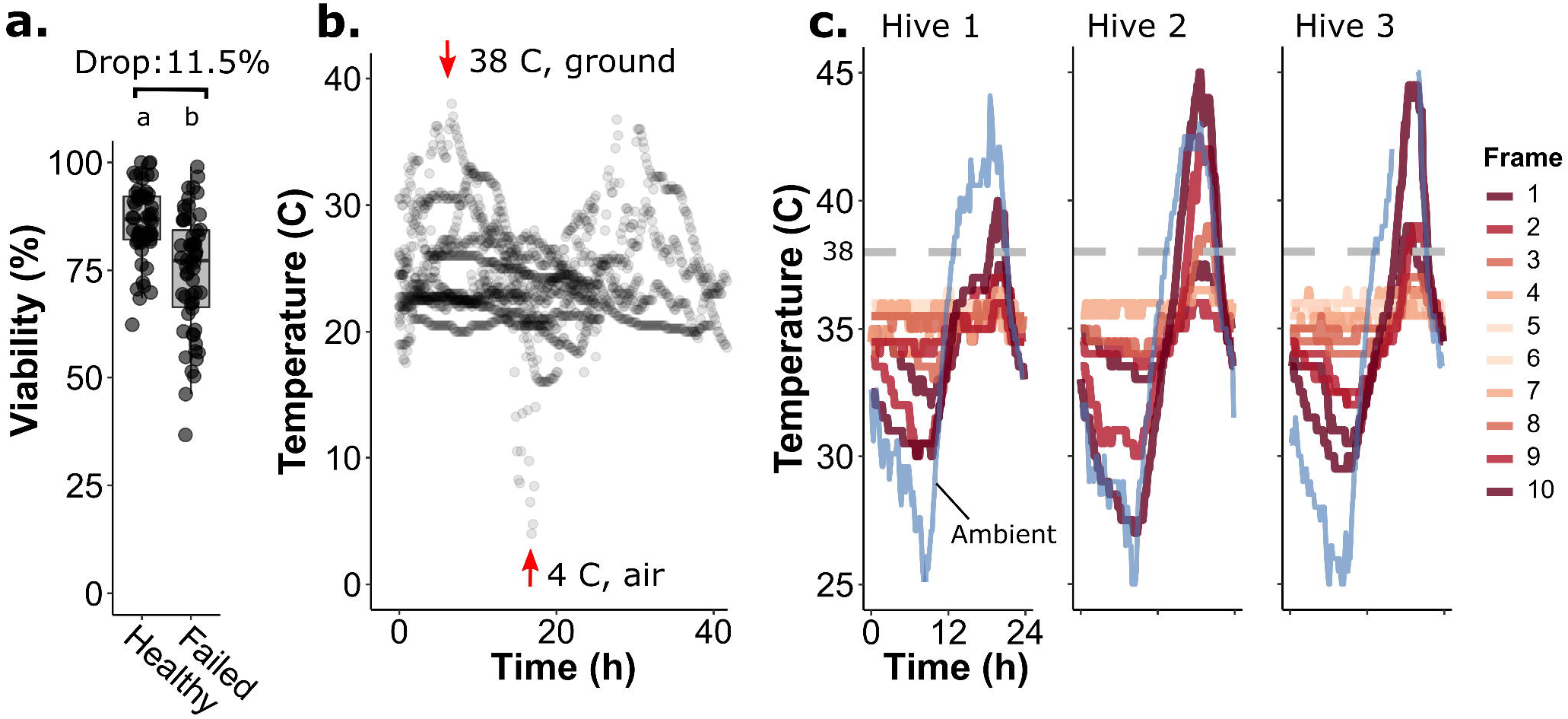
Observational data on shipping temperatures, hive temperatures, and stored sperm viability. A) Healthy (N = 55) and failed queens (N = 58) were collected from BC beekeepers for sperm viability analysis. Failed queens had significantly lower sperm viability (Student’s T-test, F = 28.16, P = 5.8×10^−6^), with an average drop of 11.5%. Boxes represent the bounds between the 2st and 3^rd^ interquartile range (IQR), midlines represent the median, and whiskers are extended by 1.5*IQR. B) Temperatures of eight domestic Canadian queen shipments were recorded during the summer of 2019 (7 via ground, 1 via air transportation). Temperature loggers were kept immediately adjacent to the queen cages. C) Internal temperatures of three standard single-deep, wooden hives were recorded, with temperature loggers placed between each frame. Ambient temperatures were recorded in the shade beneath each hive.

### Temperature spikes occur in colonies and shipments

To document if routine shipping poses a threat of adverse temperature exposure to queens, we tracked the temperatures of eight domestic queen shipments (seven via ground transportation, one via air; **Fig 1b**). We found that even in these shipments, which were not deliberately timed to occur during extreme weather events, one package experienced a temperature spike to 38°C and one dropped to 4°C. Since honey bees cannot adequately thermoregulate in queen cages, extreme ambient temperatures are a hazard for shipping. However, little is known about a whole colony’s ability to thermoregulate in the face of extreme heat.

To gain a more complete picture of temperature fluctuations within colonies, we recorded temperatures throughout the brood nest (loggers placed between each frame of three 10-frame hives) during extreme heat in August in El Centro, California. The ambient temperatures, measured in the shade beneath each hive, reached up to 45°C (**Fig 1c**). In all three hives, the temperature at the two outer-most frames spiked to upwards of 40°C for 2 - 5 h, and in two of the hives, temperatures exceeded 38°C even one or two frames closer to the core. Therefore, the colony’s ability to thermoregulate begins to break down in extreme heat, and queens can be vulnerable to temperature stress inside the hive.

### Defining critical exposure and duration thresholds

Previous research has shown that both cold (4°C) and hot (42°C) temperatures reduce stored sperm viability in queens^18^, but refined tolerance thresholds and biologically relevant viability losses are not known. To determine critical temperature and duration thresholds, we compared stored sperm viability across a temperature and duration gradient (5, 10, 15, 25, 38, 40 and 42°C, exposed for 1, 2 or 4 h followed by a 2 d recovery period) (**Fig 2a**). Not all experimental groups’ data were normally distributed (Shapiro test, P < 0.05); therefore, we analyzed it with a Kruskal-Wallis test for non-parametric data. There was a significant effect of temperature for the 2 h (⍰^2^ = 15.6, P = 0.016) and 4 h (⍰^2^ = 17.9, P = 0.0065) treatments, while not at 1 h (⍰^2^ = 9.12, P = 0.17). A Dunnett’s post hoc test revealed that the only temperatures that were significantly different from the control (25°C) were the 2 h, 10°C treatment (P = 0.045) and the 4 h, 42°C treatment (P = 0.00057), at a family-wise error rate of α = 0.05. The 2 h and 4 h data were then pooled and optimally fit to a cubic polynomial regression (R^2^ = 0.092, P = 0.012; **Fig 2b**) to find the temperature tolerance thresholds for queens, given a pre-defined maximum acceptable drop in sperm viability (11.5%). This model suggests that 15.2 – 38.2°C is the suggested “safe zone” with minimal loss of viability for 2 - 4 h exposures.

**Figure 2.**
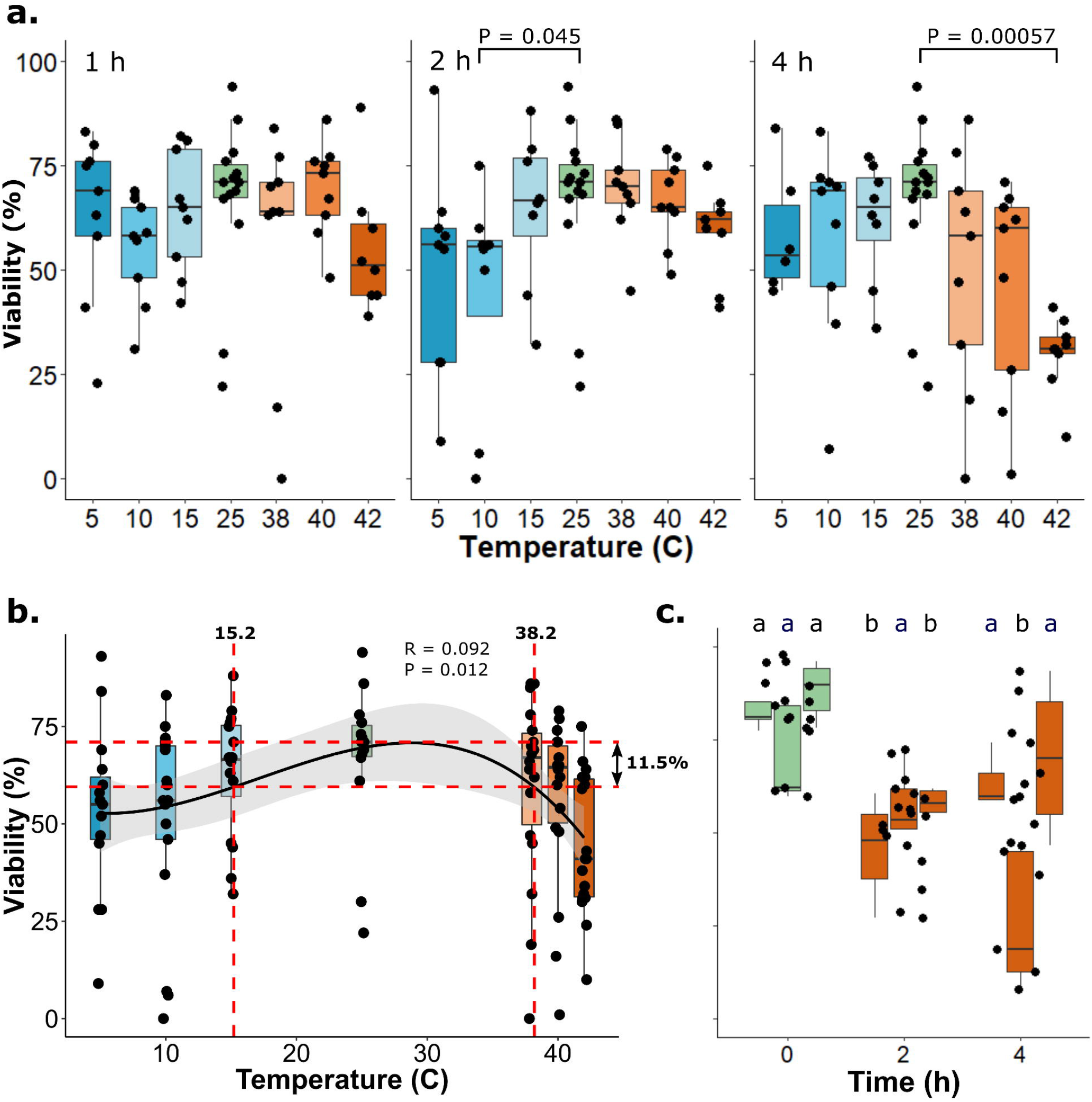
Viability of stored and ejaculated sperm after temperature stress. In all cases, boxes represent the bounds between the 2st and 3^rd^ interquartile range (IQR), midlines represent the median, and whiskers are extended by 1.5*IQR. A) Honey bee queens were heat- and cold-shocked for 1 - 4 hours, then held at 25°C prior to assessing stored sperm viability. The 25°C treatment is the negative control (the same control data is replotted in each panel). Each point represents a single queen. Data were analyzed with a Kruskal-Wallis test followed by a Dunnett’s post hoc test. See Supplementary Table 1 for sample sizes. B) Combined 2 h and 4 h treatments were optimally fit with a cubic model (R^2^ = 0.092, P = 0.012). Thresholded at 11.5% loss of viability, the line y = y_max_ – 11.5 intersects with temperatures 15.2 and 38.2°C. C) Ejaculated honey bee semen from three different colonies (5-6 drones per colony, illustrated as different boxes) was subject to heat-shock at 42°C for 2 h and 4 h, then kept at 25°C for 2 d. The total number of drones evaluated for 0, 2, and 4 h treatments were 16, 16, and 18, respectively (see Supplementary Table 2 for more details). Each data point represents a single drone. Letters indicate significant differences at P < 0.05 (two-way ANOVA, factors: colony and time, followed by a Tukey HSD post hoc test).

To test effects of heat on ejaculated sperm viability, we exposed single-drone ejaculates to 42°C for 0, 2, or 4 h, followed by a 2 d recovery period at 25°C (**Fig 2c**). These data were normally distributed (Shapiro test, P = 0.18); therefore, we used a two-way ANOVA for analysis. We found that responses differed depending on the colony source, but heat dramatically decreased viability by 35% after both 2 and 4 h (factors: time and colony, P_(time)_ = 1.2E-07, F_(time)_ = 24 df_(time)_ = 2; P_(colony)_ = 0.00015, F_(colony)_ = 11, df_(colony)_ = 2). Heat-shock therefore affects stored and ejaculated sperm viability at similar magnitudes. Drones could also be appropriate biomonitors of heat-stress, as their sperm is also sensitive to changes in temperature. However, drones are not as long-lived as queens and are only seasonally available.

### Sex biases in heat-stress survival

Sturup et al. previously reported that drones are mortally sensitive to heat^19^; however, queens are typically tolerant of stressful conditions. As a biomonitor, a favourable feature would be to survive through heat stress while accumulating a physiological and molecular record of the stress event(s). We compared drone, queen, and worker survival over time at 25°C, 38°C, and 42°C, confirming that drones, but not queens, are indeed mortally sensitive to heat (Log-rank test, P < 0.00001; Fig 3a, FigS1). Fifty-four percent of drones died over the course of 6 h at 42°C, whereas every queen survived. We also found that drones are more sensitive to heat than workers, which have a similar lifespan to drones but are non-reproductive females.

**Figure 3.**
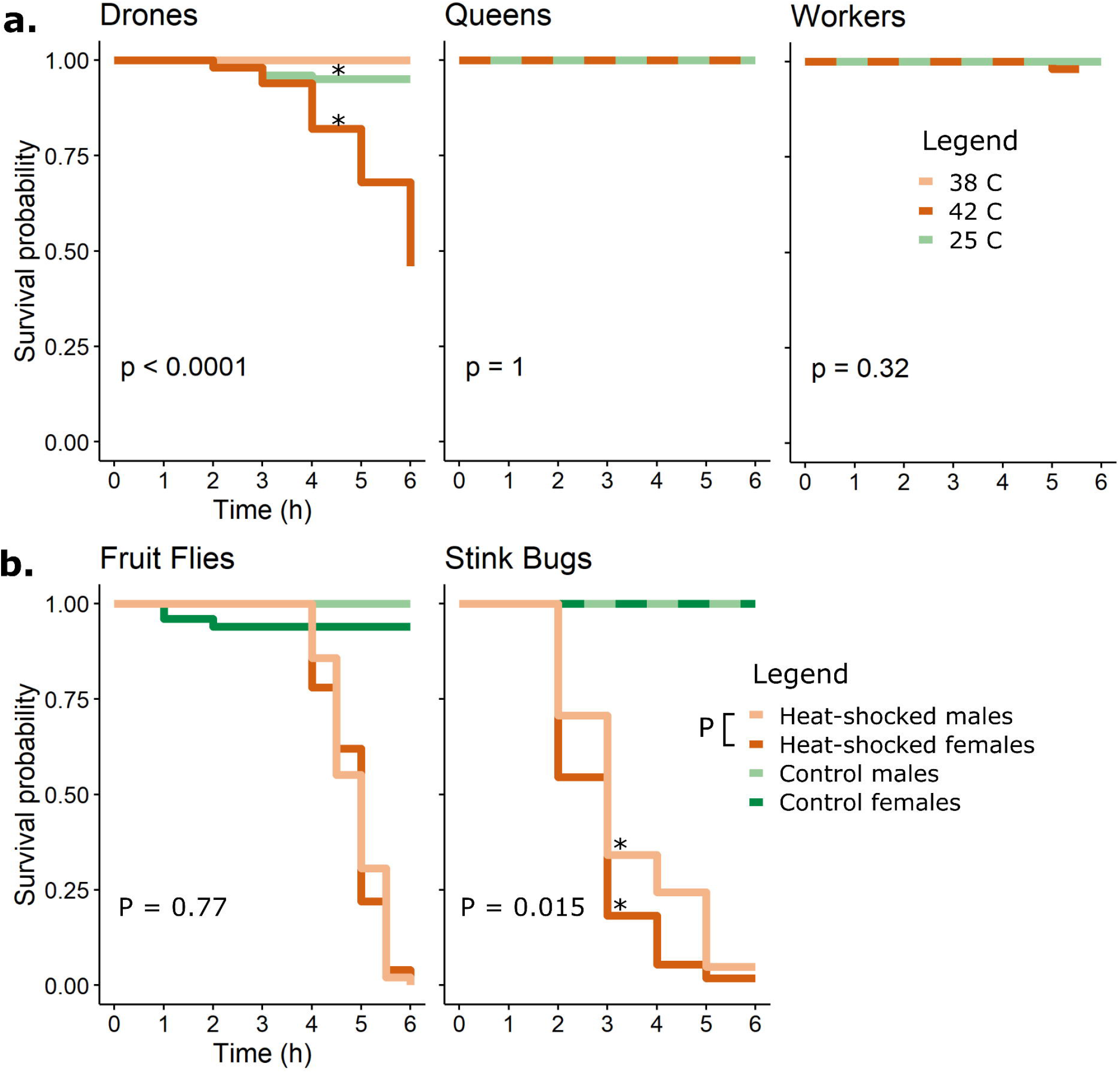
Sex-biased heat mortality in honey bees, stink bugs, and fruit flies. Statistical differences were evaluated using a log-rank test. Asterisks indicate significant differences. See Supplementary Figure 4 for risk tables, including population sizes used to generate survival curves. A) Honey bees (drones, queens, and workers) were held at different temperatures and mortality was recorded hourly. Drones are sensitive to heat, whereas queens and workers are not. B) Fruit flies (Drosophila melanogaster; heat-shock = 38°C) and stink bugs (Halyomorpha halys; heat-shock = 42°C) do not have sex-biased heat sensitivity. For fruit flies, loss of motility was used as the endpoint.

In this experiment, drones and workers were from local colonies, while queens were either from a local origin or imported from Hawaii and Australia. If bees are to be used as a global biomonitor for temperature stress, an important experiment will be to determine if there are differences in survival, physiological response, or biochemical response to heat between genetic stocks that may be adapted to hotter or cooler climates.

Honey bees’ sex-biased heat sensitivity is puzzling because drones and queens spend most of their lives inside the hive and can avoid participating in mating flights during hot weather. Therefore, they are exposed to similar environmental conditions, and based on this, one might expect them to have similar physiological tolerance thresholds. However, the ability of a colony to survive and produce reproductive individuals directly depends on the survival of individual queens (there is only one per colony), and not individual drones (hundreds per colony). Natural selection may have favoured high survivorship of queens, as evidenced by not only their long lifespan (up to 5 y) but also their high tolerance to heat stress, despite the death of their stored sperm. To our knowledge, sex biases in thermal tolerance for other social insects has not yet been investigated, as critical thermal tolerance studies have focused on workers or on male fertility alone.^46–55^

Next, we investigated sex biases in heat tolerance for two solitary insect species—fruit flies (*Drosophila melanogaster*) and brown marmorated stink bugs (*Halyomorpha halys*). The stink bugs have a similar body size and thermal tolerance range as honey bees^56^ and their global distribution has expanded to cover four continents, making them a potential candidate biomonitor. Fruit flies have a much smaller body size but are also widely distributed, so they could be candidate biomonitors too. We found that both male and female stink bugs and fruit flies readily die with heat (**Fig 3b and c**), meaning that they cannot accumulate a physiological or molecular record of a heat stress event and have limited utility for observing impacts of heat on fertility through time. However, these data do indicate another worrying trend: that exceptionally extreme temperatures may reduce insect populations through direct kills, in addition to reducing reproductive output if they survive. We found no difference between males’ and females’ heat sensitivities for fruit flies and a slight opposite (female) biased heat sensitivity in stink bugs (P = 0.015, log rank test). In contrast, the high survivorship of honey bee queens makes them a good candidate biomonitor, should they possess molecular signatures indicative of heat stress when it occurs.

### Heat activates specific proteins in the spermatheca

Sperm longevity is enabled by molecular processes that reduce oxidative damage and maintain sperm in a quiescent metabolic state. For example, the spermatheca is a highly anaerobic environment, which helps prevent reactive oxygen species (ROS) formation^57^. Enzymes that further limit damage from ROS are also upregulated in mated queens compared to virgins^58,59^, and ROS damage leads to infertility in mammals^60,61^. Heat is well known to lead to oxidative stress^62^; therefore, we hypothesized that queens may combat heat-stress by upregulating enzymes that mitigate oxidative damage. Additionally, glyceraldehyde-3-phosphate dehydrogenase (GAPDH) has been previously implicated in stored sperm longevity, since metabolic activity measurements of stored sperm supplemented with its substrate (glyceraldehyde-3-phosphate) improved viability and produced the highest molar ratio of ATP via anaerobic catabolism^57^. Heat-induced changes in GAPDH expression could therefore also impact sperm viability.

To test these hypotheses, we compared expression of ROS mitigating enzymes (superoxide dismutases (SOD1, 2, and 3) and catalase) as well as GAPDH in heat-shocked and non-heat-shocked virgin spermathecae, mated spermathecae, and ejaculated semen. Stored sperm are reportedly transcriptionally active in queens to some degree^57^, although whether this is true in general has been a matter of debate across species^63^. It is not known with certainty whether ejaculated sperm are transcriptionally active; nevertheless, analyzing ejaculates, virgin spermathecae and mated spermathecae helped us to disentangle the male versus female origins of expression in sperm-filled, mated spermathecae. We found that heat-shock did not upregulate expression of ROS-mitigating enzymes, nor GAPDH (**Supplementary Figure 1**); rather, all enzymes were consistently (but not significantly) downregulated with heat (by 10-30%). Queens must therefore employ other strategies, if any, to combat heat stress.

A multitude of other proteins could also be responsible for mitigating damage from heat—most obviously, members of the heat-shock protein (HSP) family^24,64–66^. HSPs generally function as molecular chaperones that stabilize proteins, refold damaged proteins, and prevent protein aggregation, but they can have diverse functions in specific contexts and their precise role in the honey bee spermatheca is unknown. Some HSPs contain an ATPase domain and require ATP to function (*e.g.,* HSP70s, HSP90s, HSP110s), while others operate independently of ATP (*e.g.,* HSP10s, HSP20s, HSP60s). In addition, since glyceraldehyde-3-phosphate yields the most ATP per unit via anaerobic catabolism of any tested substrate and significantly increased sperm longevity^57^, this implies that ATP usage economy is a critical factor for maintaining viable sperm. Therefore, we expect that ATP-independent HSPs should be upregulated with heat stress in spermathecae, and ATP-dependent HSPs (or a mix of ATP-dependent and ATP-independent HSPs) should be upregulated as a result of heat stress in other tissues. In addition, some ATP-independent short HSPs (sHSPs, typically 20-28 kDa) suppress ROS generation while preventing protein aggregates and apoptosis^62,67^. Since molecular processes that reduce oxidative damage favour sperm longevity, we hypothesized that heat should induce expression of sHSPs in the spermatheca.

To determine which HSPs were upregulated with heat-shock, we compared global protein expression profiles in heat-shocked and non-heat-shocked mated and virgin spermathecae, ejaculates, and ovaries (**Fig 4a, Supplementary Figure 2**). We hypothesized that the heat-shock response should both mitigate ROS production and conserve ATP in the spermatheca, but this ATP-conservation may not be as critical in the ovaries where the ATP economy is not expected to be as tightly controlled. Of the 2,778 protein groups identified in the spermathecae and ejaculates, only five were significantly up-regulated with heat (5% FDR), all of which were identified in the mated and virgin spermathecae. All five of the proteins were unique, ATP-independent sHSPs (accessions: XP_001120194.1, XP_001119884.1, XP_395659.1, XP_001120006.2, and XP_026294937.1) (**Fig 4b**). In heat-shocked virgin spermathecae, two of the same sHSPs (XP_395659.1 and XP_001120006.2) were also significantly upregulated, and all mirrored the expression patterns in the spermathecae of heat-shocked mated queen (even if not significant), indicating that this is a queen-derived, and not a sperm-derived, response (**Fig 4c and d**). By contrast, the most strongly upregulated protein in heat-shocked ovaries was HSP70—an ATP-dependent HSP— and no sHSPs were upregulated in this tissue, supporting our initial hypotheses (**Fig 4e**).

**Figure 4.**
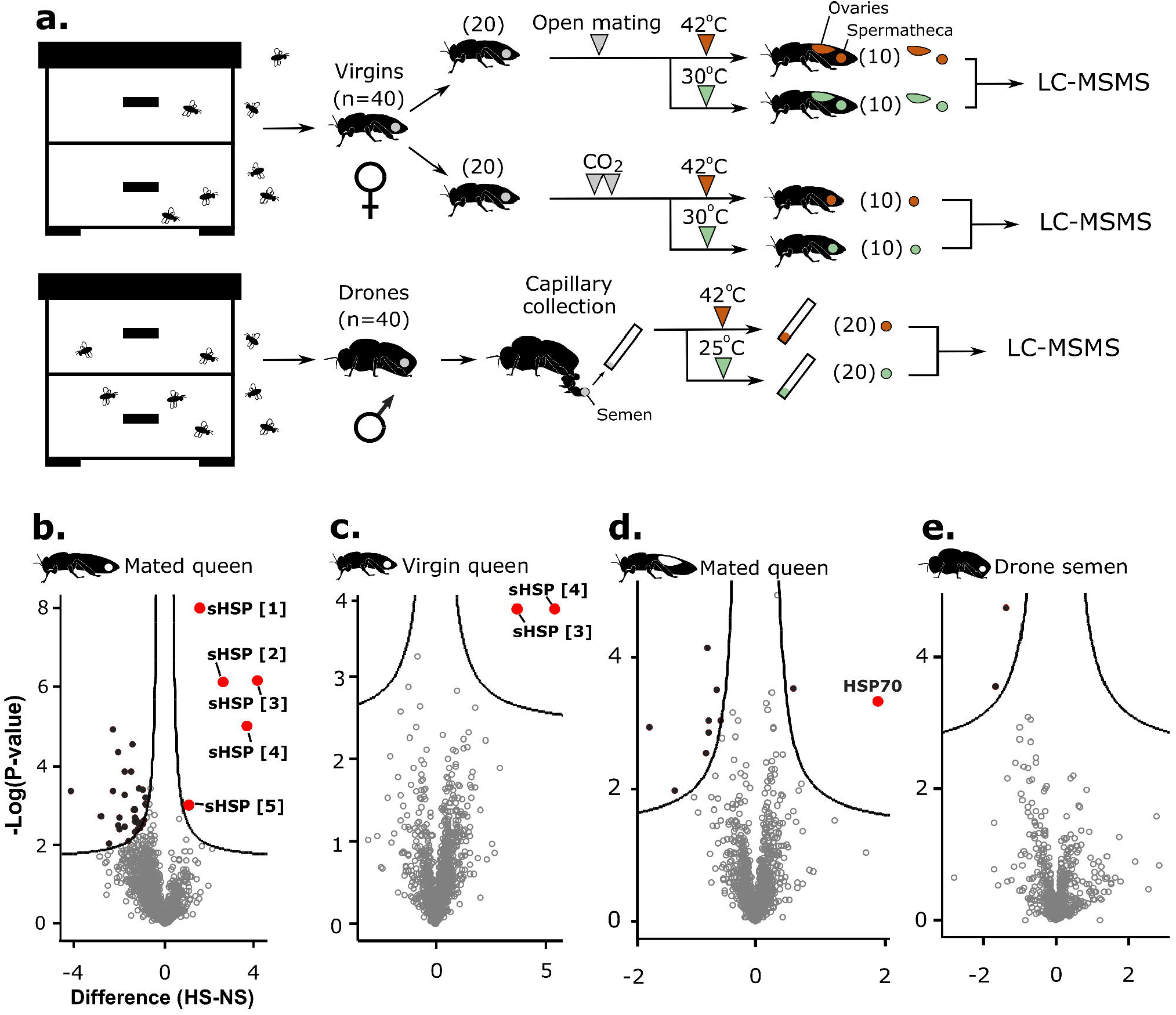
Differential protein expression comparing heat-shocked (HS) and not heat-shocked (NS) reproductive tissues. A) Samples included mated queen spermathecae, virgin queen spermathecae, ejaculated semen, and mated queen ovaries, which were all analyzed by intensity-based, label-free quantitative tandem mass spectrometry. For drones, 4 semen samples were pooled into one replicate, then fractionated into eight fractions by basic reverse phase chromatography prior to analyzing by mass spectrometry (final biological replicates: n = 5 heat-shocked and 5 non-heat-shocked). The significance cut-off for volcano plots (B-E) was 10% FDR (false discovery rate, permutation-based). Volcano plots represent mated queen spermathecae (B), virgin spermathecae (C), mated queen ovaries (D), and drone semen (E). Differentially expressed HSPs are red; other differentially expressed proteins are black. Accessions for sHSP [1-5] are XP_001120194.1, XP_001119884.1, XP_395659.1, XP_001120006.2, and XP_026294937.1, respectively. The accession for HSP70 is NP_001153536.1. Bee silhouettes are adapted from McAfee et al. (2019)^84^ (CC-BY 4.0).

As expected, the significantly enriched Gene Ontology (GO) terms were all related to responses to heat and stress in the spermathecae (**Fig 5a and b**). However, the only significantly enriched GO term in the semen analysis was related to the electron transfer activity (**Fig 5c**), driven by a heat-induced downregulation of the proteins linked to this GO term (**Fig 5d**), suggesting that the sperm may be less able to produce the large amounts of ATP necessary for flagellar beating. This is consistent with the findings of Gong *et al.,* who found that heat stress at 42°C impaired mitochondrial function, reduced electron transport chain complexes’ activities, and lowered total cellular ATP^23^. Numerous proteins (none of which were HSPs) were downregulated in heat-shocked spermathecae, but based on the GO term enrichment analysis and manual inspection of their functions, it is unclear what their biological significance is. It is possible that they are degradation products of heat-killed sperm, but since these proteins were largely absent in the semen samples, we cannot confirm this hypothesis.

**Figure 5.**
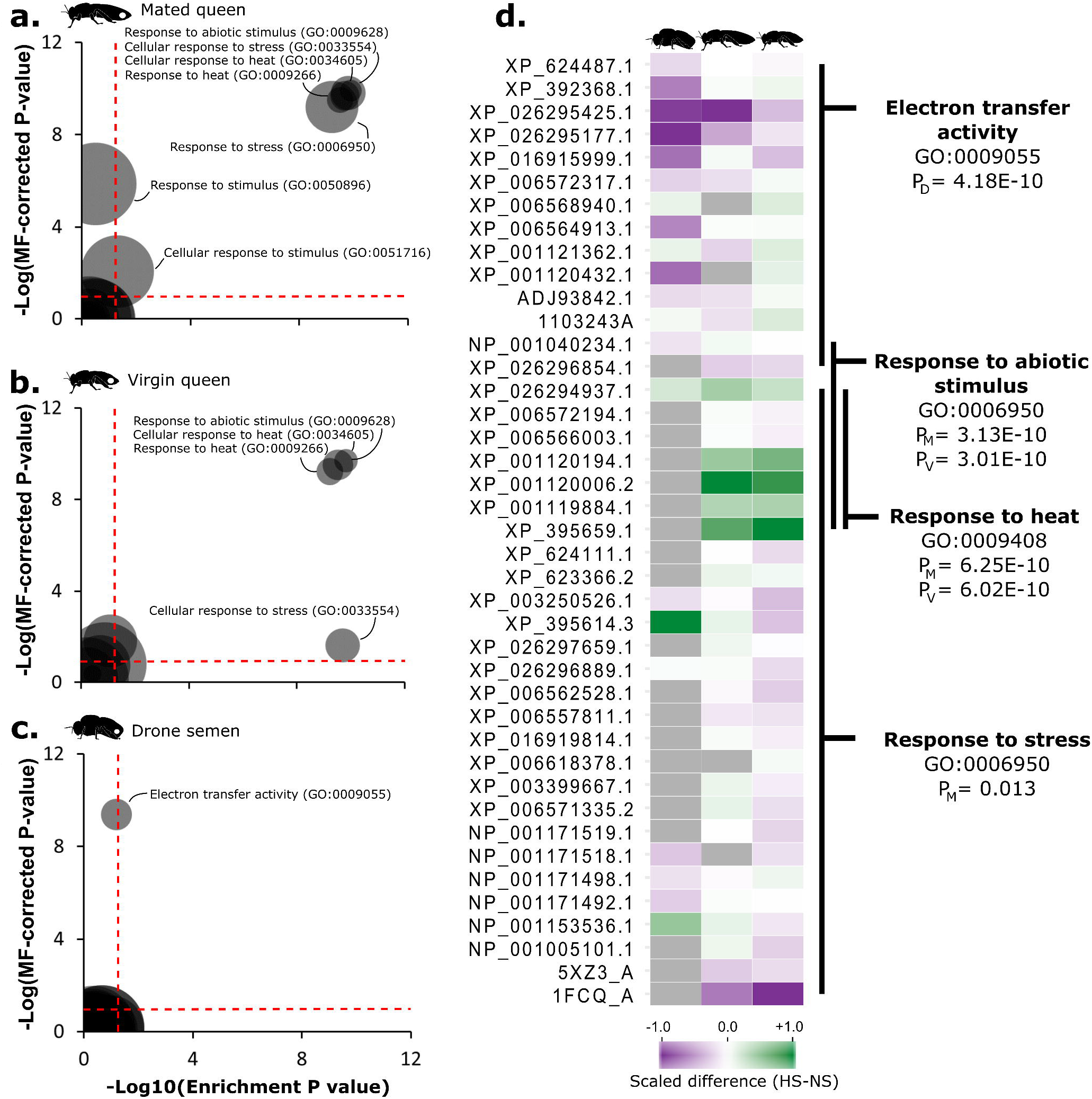
Gene ontology (GO) term enrichment analyses. A-C) Enrichments were performed via gene score resampling, correcting for both multiple hypothesis testing and protein multifunctionality (MF). The x-axes depict enrichment P values (10% FDR, Benjamini-Hochberg correction) without multifunctionality correction. The y-axes depict enrichment P values after correction for both multiple hypotheses and multifunctionality. The red dotted lines indicate the P value significance cut-off that achieves 10% FDR. D) Summary of proteins associated with significantly enriched GO terms. Subscripts refer to comparisons within drones (D), mated queens (M), and virgin queens (V). Bee silhouettes are adapted from McAfee et al. (2019)^84^ (CC-BY 4.0).

Twenty HSPs were identified in spermathecae overall and 13 were identified in the semen (**Fig 6**); however, the precise functions of specific honey bee HSPs are largely unknown. To gain insight into potential roles of the five differentially expressed HSPs we identified, we compared their sequences to annotated sequences in other species and identified putative protein domains using NCBI’s basic local alignment search tool (BLAST)^68^. All five of the HSPs contain one or more alpha crystallin domains, which is characteristic of small HSPs (sHSPs). Four of these HSPs are within the expected molecular weight range, and one of the proteins (XP_026294937.1) is predicted to be 56.7 kDa (and contains two alpha crystallin domains instead of one), highlighting that HSPs should not necessarily be categorized based on molecular weight alone. All five of the honey bee sHSPs are paralogous to the *D. melanogaster* gene *l(2)efl* (also known as CRYAB). In *Drosophila*, upregulation of this gene causes increased lifespan of individual flies^69^. CRYAB and other sHSPs are highly conserved in both vertebrates and invertebrates,^70^ but is by far the best studied in human. In humans, sHSP upregulation is associated with anti-apoptotic properties, as well as mitigating ROS production^62,66,71,72^. Their up-regulation in heat-shocked testes is thought to help compensate for the damaging effects of heat^71^, and we speculate that they are playing a similar role in the spermathecae. Queens with strongly up-regulated sHSPs may therefore be better able to counter-act ROS production, sperm death, and ultimately maintain longevity. However, it remains unknown if the honey bee paralogs have functionally diverged from those in other species.

**Figure 6.**
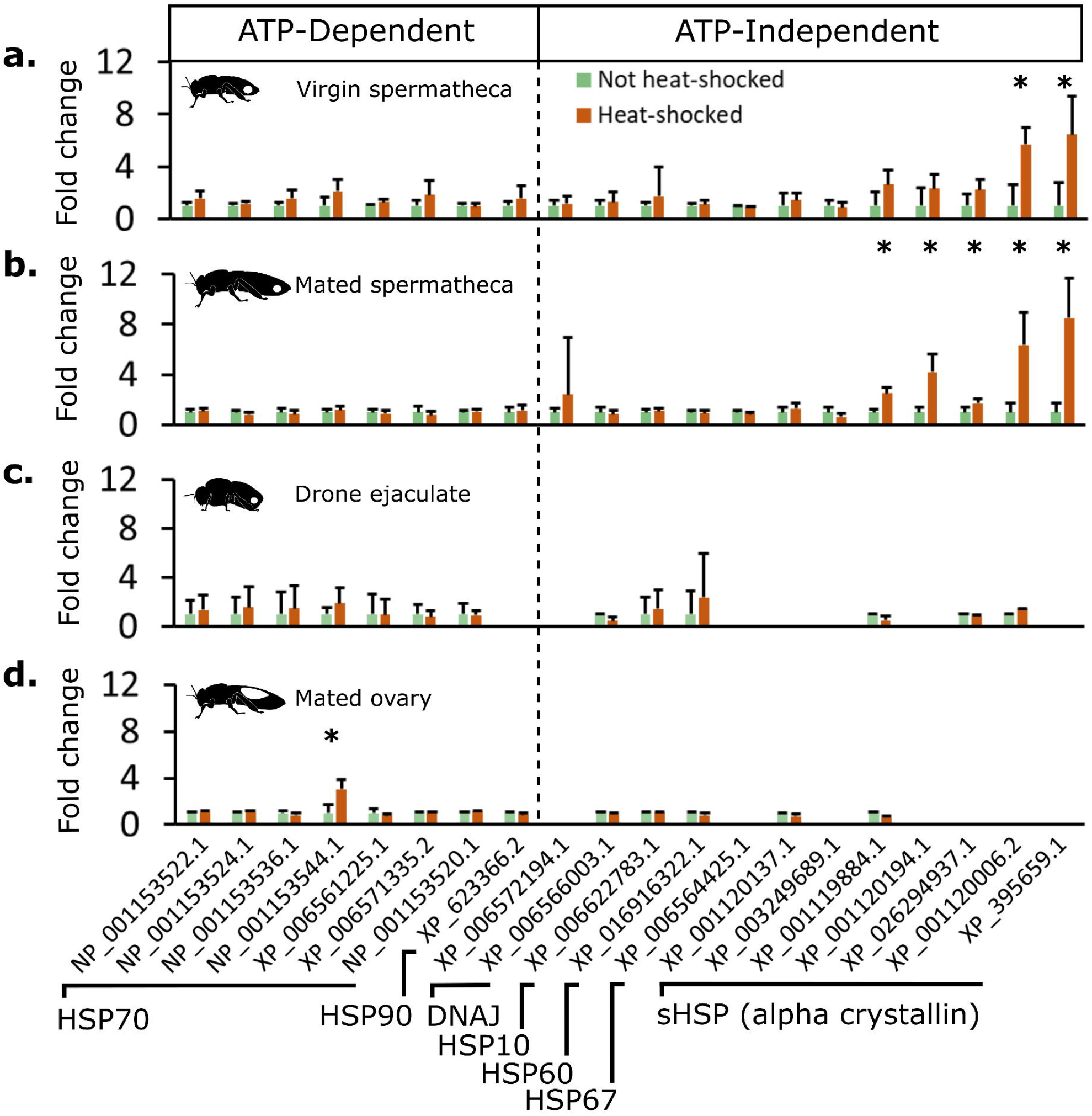
Heat-shock protein expression profiles. Proteins with GO terms for cellular response to heat were retrieved from the proteomics datasets. Fold-change is relative to the non-heat-shocked state. Proteins marked with an asterisk were significantly up-regulated in the global differential gene expression analysis (permutation-based FDR: 10%), and error bars indicate standard deviations.

Other researchers have suggested that in mammals, some ATP-dependent HSPs (e.g., HSP70s and HSP90s) may be important for maintaining male fertility via quality control of sperm^66,73^. Unlike sHSPs, these HSPs appear to be pro-apoptotic factors and could theoretically help prevent damaged sperm from being able to fertilize an egg. Neither queen spermathecae nor drone ejaculates provide evidence supporting this strategy of quality control in honey bees; in this experiment, none of the HSPs that were up-regulated with heat in spermathecae or semen contained an ATP-binding nor ATPase domain, which are characteristics of HSP70s and HSP90s but not the sHSPs (**Figure 5 and 6**)^66^. Rather, HSP70 was only upregulated in the ovaries, which does not directly participate in sperm maintenance.

In mammals, sHSPs are upregulated in testes after heating, but so are ATP-consuming proteins like HSP105, HSP70-1, HSP70-2, and HSP90^74–76^. We speculate that the energetic cost associated with ensuring high sperm quality is advantageous in mammals because it helps reduce the risk of an egg going unfertilized, which would produce no progeny. However, the honey bee’s differing strategy of upregulating only the ATP-independent sHSPs in the spermatheca is consistent with a tightly controlled ATP usage economy, as has been previously suggested^37,57^. The ATP-independent sHSPs have the dual purpose of also limiting oxidative damage and conserving ATP. The significant enrichment for differentially expressed proteins involved in multiple nucleotide metabolic processes, including ATP, in the transition from virgin to mated spermathecae (**Supplementary Figure 2**) supports the notion that regulating ATP production and consumption is critical for maintaining stored sperm viability.

Moreover, analyzing heat-shocked mated queen ovaries revealed that HSP70—an ATP-dependent HSP— was uniquely upregulated in this tissue (**Fig 5 and 6**). This observation is consistent with selection for ATP-consuming quality control mechanisms when a failed fertilization event fails to produce offspring (in this case, when a non-viable egg meets a viable sperm). Overall, these data indicate that upregulation of specific sHSPs is not a general indicator of cellular stress, since it was not observed in the ovaries. Indeed, the ovaries appear not to express most sHSPs at all, whereas they were abundantly expressed even in non-heat-shocked spermathecae.

Finally, we compared protein expression in spermathecae, fat bodies, and ovaries between 11 queens that failed due to unknown causes and 11 age-matched, apiary-matched healthy queens. We identified 1,219, 1,640, and 1,782 proteins, respectively, but did not find any significant expression differences at 10% FDR, indicating that there is not one universal signature of failure (**Supplementary Figure 3**). Rather, we suspect that different stressors alter protein expression in these tissues in different ways, and that no significant differences were found because it is highly unlikely that the queens all failed from the same cause.

We propose that the sHSP response of the spermathecae may serve as a post-queen-failure biomarker of heat stress which could help diagnose causes of colony failure in the field and simultaneously provide an indication of risks for heat-induced loss of fertility for other species. Beekeepers routinely replace queens every 1-2 y, and these queens could be banked for biomarker testing as part of a surveillance program. Future experiments should include a blind heat-shock trial to determine a) if previously heatshocked queens can be reliably distinguished from non-heat-shocked queens based on these biomarkers, b) how long the heat-shock proteomic signature lasts, and c) if other stressors produce proteomic signatures that overlap with the heat-shock signature.

## Conclusion

Our experiments show that temperature stress deeply damages both stored and ejaculated honey bee sperm viability, that queens are vulnerable to temperature changes both in colonies and during transport, and that temperatures ranging between 15°C and 38°C for 2 - 4 h are generally safe for queens. Honey bees have a strong sex bias in heat tolerance, with females being highly tolerant—a bias which does not exist in the two solitary insect species we tested (*H. halys* and *D. melanogaster*). Future research should investigate if this sex-biased heat tolerance is also present in other Hymenopterans in order to better understand the evolutionary origin. Upon heat-shock, queens upregulate ATP-independent HSPs in their spermathecae, which both minimizes ATP consumption and could provide beneficial anti-apoptotic properties. In contrast, HSP70 (an ATP-consuming HSP) is upregulated in ovaries. Once validated in field trials, these protein signatures could serve as biomarkers for heat stress enabling longitudinal surveys for the prevalence of heat-induced loss of sperm viability in diverse landscapes as part of a biomonitoring program.

## Methods

### Sperm viability assays

Honey bee queens (Kona Queens supplied in a single shipment) were treated at one of five different temperatures (5, 10, 15, 25, 38, 40, and 42°C) for 1, 2, or 4 hours, then held at 25°C for 2 d. The temperature range was chosen because previous research showed that 4°C and 8°C were sufficiently cold to reduce sperm viability, and that 40 - 42°C was sufficiently hot.^18^ Therefore, more temperatures were chosen at the cold (5, 10, and 15°C) and hot (38, 40, and 42°C) extremes to try to capture the critical point at which the viability begins to drop. We chose 25°C as the control temperature, rather than 35°C, because these queens were obtained from a commercial supplier and queens are not held at hive temperature during transport. While a 35°C treatment group would be an appropriate control, it would not be meaningful in these circumstances.

Fourteen queens (replicates) were included in the 25°C treatment (negative control), whereas 8 or 9 queens were included in all other temperatures and exposure durations (see **Supplementary Table 1** for specific replication information). Following this, queens were beheaded, and their spermathecae were dissected with fine forceps. The spermathecae were gently agitated with a pipet tip in 100 μl Buffer D (17 mM D-glucose, 54 mM KCl, 25 mM NaHCO_3_, 83 mM Na_3_C_6_H_5_O_7_) to break them open and release the sperm. Sperm viability was determined using a live/dead sperm viability kit (Thermo) following the protocols of Collins and Donoghue^77^. Briefly, the SYBR14 dye was diluted 1:9 in DMSO. Two microlitres of the diluted SYBR14 dye and 4 μl of propidium iodide were gently mixed with the sperm and incubated for 15 minutes in the dark. Two microlitres of the mixture was then added to a glass microscope slide and viewed under a fluorescent microscope. Live (green) and red (dead) sperm were counted until 100 cells were observed, covering multiple fields of view. Unless otherwise reported, all statistical analyses were performed in R (v3.5.1). First, the data were tested for normality using a Shapiro test. The data was not normally distributed, we used a non-parametric test (Kruskal Wallis) to investigate the effects of temperature on viability for each duration separately. When a significant effect of temperature was identified, we performed a post-hoc Dunnett’s test to identify significant contrasts to the 25°C control. To identify the best-fitting linear model, we pooled the 2 h and 4 h data and performed linear regressions using temperature as a continuous variable, testing exponents 1 - 4 to identify the optimal fit (highest R^2^).

Drones were harvested from three different colonies headed by Kona queens with unknown relatedness. The colonies were kept in Beaverlodge, Alberta. Semen was collected with glass capillaries according to the methods of Collins and Donoghue^77^. Briefly, we pinched drone abdomens with a rolling motion from the anterior to posterior end to expel the endophallus and semen. The semen was collected by first filling glass capillaries with 1 μl Buffer D, then drawing up the semen via capillary action (avoiding the white mucus secretions). Capillaries were then filled with a further 1 μl of Buffer D to prevent the sample from drying out, and both ends were wrapped with parafilm. This technique typically yielded 0.5-1.0 μl of semen per sample. Semen samples were then heat-shocked at 42°C for 0, 2, or 4 h, then allowed to recover at 25°C for 2 d. Depending on the colony, 5 or 6 drones were sampled for each time point (see **Supplementary Table 2** for complete replication information). We chose a lower temperature for semen recovery because ejaculated drone semen can be maintained at room temperature for several weeks while retaining high viability^78^. Viability assays for these semen samples was performed following the same methods as for spermathecae. Data were analyzed by a Shapiro test and Levene’s median test to confirm normal distribution and evaluate the validity of the assumption of equal variance, respectively. The data passed both tests, so a two-way ANOVA (factors: time, colony) was performed, followed by a Tukey HSD test.

### Recording temperatures during shipments

We arranged seven shipments of locally produced (BC) queens via ground transportation from Kettle Valley Queens, Nicola Valley Honey, Wild Antho, Campells Gold Honey, Heather Meadows Honey Farm, Six Legs Good Apiaries, and Wildwood Queens (located in Grand Forks, Merrit, Armstrong, Abbotsford, Telka, Surrey, and Powell River, respectively) to the University of British Columbia, Vancouver, in the summer of 2019. These queens (7-8 queens per shipment) were all considered to be “good” queens by the donating bee breeders based on their laying pattern. These queens cumulatively make up the “healthy” queen cohort (N = 55 queens total). An additional shipment travelled from Edmonton to UBC Vancouver via air freight. All shipments were accompanied by two temperature data loggers (B-series WatchDog loggers, Spectrum Technologies) set to record the temperature every 10 minutes for the duration of the shipment.

### Failed queen viability survey

We solicited samples of failed queens, as rated by beekeepers, from throughout BC in the summer of 2019. Reported reasons for designating the queens as ‘failed’ included poor brood pattern, drone laying, poor population build-up, disease symptoms, injury, small size, and worker rejection. These queens (N = 58) make up the “failed” queen cohort. We measured the sperm viability of all failed and healthy queens using the same live/dead sperm staining kit as described under “Sperm Viability Assays.” These queens were not age-matched. Therefore, the tolerance threshold loss of viability we set using these queens is likely a conservative estimate.

### Hive temperature recording

To record the internal and external temperatures of colonies, we placed temperature loggers inside three standard, 10-frame colonies in El Centro, California, from August 23^rd^ to September 6^th^, 2017. The hottest day (the data shown) was August 26^th^. Loggers were set to record the temperature every 10 minutes and were placed between each frame of the hive. Hives consisted of 10 deep frames in a single wooden Langstroth brood box with a medium depth box on top of the brood chamber and a migratory wooden cover. The ambient temperatures were recorded with a temperature logger placed in the shade beneath each hive.

### Heat-shock survival

For honey bee heat-shock tests, drones and workers were collected by retrieving a frame of capped brood from a hive and allowing the bees to emerge in an incubator. Newly emerged bees were marked with a paint pen and returned to the colony to age for one week, at which time they were recaptured and caged in California mini cages. For drone heat-shock tests, one drone was added per cage along with malleable candy (icing sugar-based) and five worker attendants. A total of 199 drones were caged in the following experimental groups: 42°C heat (50), 38°C heat (50), and 25°C (99) for six hours (all at 60% relative humidity). Every hour, the number of drones that perished was recorded. Worker heat stress tests were conducted independently with six workers per cage (54 workers per group) held at 42°C and 25°C.

Queens were from a variety of sources (local, Hawaii, and Australia) and their ages ranged from approximately 3 - 5 weeks. They were kept in California mini-cages with five worker attendants each and held at 42°C, 38°C, or 25°C for either 1, 2, or 4 h as part of other experiments (for viability measurements and proteomics). Relative humidity varied between 40 and 80%, but this did not have an effect of survival (100% of queens survived in both cases). Eight queens were held at 42°C for 6 h at a constant 60% relative humidity specifically for survival analysis. See **Supplementary Figure 4** for sample sizes and risk tables for each treatment group. No queens died in these tests; therefore it was not necessary to account for geographic origin in statistical analyses.

Male and female 3- to 10-day-old Canton S wild type fruit flies were held at 38°C and 25°C for six hours in *Drosophila* vials with media (50 males and 50 females). Loss of motility was recorded every hour. We used this as the endpoint because fruit flies will become paralyzed with heat, but recover motility after several hours (in some cases, overnight) at cooler temperatures. Control female flies were maintained for a further week after the experiment to confirm that the majority of them had successfully mated prior to the heat stress tests.

Male and female stink bugs were reared in the laboratory according to standard protocols^79^. Stink bugs which transitioned from nymphs to adults 3 - 7 d prior were transferred to ventilated 500 ml volume plastic cages with a piece of moist paper towel (3 - 6 stink bugs per cage). They were heat stressed at 42°C (55 females and 41 males) or held at 25°C (36 females and 24 males) for 6 h. Only mated females carrying eggs were included in the experiment, as determined by post-stress dissection.

In all cases, Kaplan-Meier survival curves were generated in R and compared using log-rank tests. See **Supplementary Figure 3** for risk tables and sample size information for survival tests for all species.

### Heat-shock for proteomics

Our experimental design for the proteomics experiments was to compare heat-shocked and non heatshocked treatments of three tissues: virgin spermathecae, mated spermathecae, and ejaculated semen (N = 10 each for all but the semen, which was N = 5 where each N was a pooled sample of five drones from one colony). For the mated spermathecae, expression differences induced by heat-shock would be the sum of any expression differences that may have occurred in the queen’s cells and in the sperm cells. The virgin spermathecae, however, do not contain sperm so that response to heat-shock is purely from the queen.

Honey bee colonies were maintained in an apiary at the University of British Columbia. During the summer of 2018, 40 queens were reared from a single colony of local origin and half were allowed to open mate, while the other half were kept as virgins in plastic queen cages. Two weeks after emergence, the virgin queens were given two, eight-minute carbon dioxide treatments on sequential days, then reintroduced to their nucleus colonies. This process stimulates virgin queens to begin laying^80^, and we conducted these treatments in order to minimize the physiological differences between virgin and mated queens.

Virgin and mated queens were retrieved from their nucleus colonies and half of each (10) were subjected to heat-shock (42°C, 2 h), and then maintained at 30°C for 2 d. The other half were held only at 30°C for 2 d. Four to six weeks after mating, the queens were anesthetized with carbon dioxide, beheaded, then their spermathecae were removed with fine forceps. Both ovaries were also removed and weighed.

During the same summer, 200 drones from a different colony in the same apiary were collected and maintained in the laboratory overnight at ambient temperature with excess syrup (50% sucrose). The next day, semen was harvested with glass capillaries according to the methods described above. Because many drones were not sexually mature, 60 semen samples (out of the 200 drones) were collected. Capillaries were placed in petri dishes and half (30) were heat-shocked as described above, then kept at 25°C for 2 d. The other half were only kept at 25°C for 2 d. Ten samples from each experimental group were used for sperm viability assays as described above.

### Proteomics sample preparation

Semen and spermatheca samples were homogenized in 2 ml screw-cap tubes containing 100 μl of lysis buffer (6 M guanidinium chloride, 100 mM Tris, pH 8.5) and four ceramic beads. The homogenizer (Precellys 24, Bertin Instruments) was set to 6,500 s^−1^ for 30 s, then samples were centrifuged (16,000 rcf, 10 min, 4°C) to remove debris. Supernatants were transferred to a new tube and diluted 1:1 with dH_2_O. Protein was precipitated by adding four volumes of ice-cold acetone and incubating overnight at - 20°C. The precipitated protein was pelleted and washed twice with 500 μl of 80% acetone, then the pellet was allowed to air dry (ca. 5 min) prior to solubilization in 50 μl of digestion buffer (6 M urea, 2 M thiourea).

Approximately 25 micrograms of protein were reduced (0.5 μg dithiothreitol, 20 min), alkylated (2.5 μg iodoacetamide, 30 min, dark), and digested (0.5 μg Lys-C for 3 h, then 0.5 μg trypsin overnight). Digested peptides were acidified with one volume of 1% trifluoroacetic acid and desalted with high-capacity STAGE tips as previously described^81^. Eluted samples were dried (SpeedVac, Eppendorf, 45 min) and resuspended in Buffer A (0.1% formic acid). Peptide concentrations were determined using a NanoDrop (Thermo, 280 nm) and sample orders were randomized for liquid chromatography-tandem mass spectrometry (LC-MS/MS) analysis.

### LC-MS/MS data acquisition and analysis

Peptides (0.5 μg for each sample) were injected on an EASY-nLC 1000 liquid chromatography system (Thermo) coupled to an Impact II Q-TOF mass spectrometer (Bruker), essentially as previously described^82^. The LC system included a fused-silica (5 μm Aqua C18 particles (Phenomenex)) fritted 2 cm trap column connected to a 50 cm analytical column packed with ReproSil C18 (3 μm C18 particles (Dr. Maisch)). The separation gradient ran from 5% to 35% Buffer B (80% acetonitrile, 0.1% formic acid) over 90 min, followed by a 15 min wash at 95% Buffer B (flow rate: 250 μL/min). The instrument parameters were: scan from 150 to 2200 m/z, 100 μs transient time, 10 μs prepulse storage, 7 eV collision energy, 1500 Vpp Collision RF, a +2 default charge state, 18 Hz spectral acquisition rate, 3.0 s cycle time, and the intensity threshold was 250 counts.

Mass spectrometry data were searched using MaxQuant (v1.5.3.30) using default parameters, except “match between runs” was enabled. Peptide spectral matches, peptide identifications and protein identifications were controlled at 1% false discovery rates (FDRs). The protein search database was the NCBI Identical Protein Groups database for **Apis mellifera** (downloaded Nov. 1^st^, 2018; 21,425 entries) plus all honey bee viral proteins contained within NCBI (a further 508 entries). Differential expression analysis was performed in Perseus (v1.6.1.1) essentially as previously described^82^. Histograms of protein counts across LFQ intensities were first inspected for normality. Then proteins differentially expressed between heat-shocked and non-heat-shocked samples, as well as among tissues (semen, virgin spermathecae, and mated spermathecae), were identified using two-tailed t-tests and an ANOVA, respectively. Only P values surviving the 10% FDR threshold (permutation-based method) were considered significant.

### GO term enrichment analysis

Gene Ontology (GO) terms were retrieved using BLAST2GO (v4.1.9)and subsequent enrichment analyses were conducted using ErmineJ^83^. We used the gene score resampling (GSR) method (with P values as scores). Unlike conventional over-representation analyses, this method does not depend on submitting a ‘hit list,’ the composition of which is sensitive to arbitrary significance cut-offs. Rather, the GSR method uses p-values as a continuous variable and looks for GO terms that are enriched in proteins with low p-values along the continuum. More documentation about the GSR method and ErmineJ can be found at https://erminej.msl.ubc.ca/help/tutorials/running-an-analysis-resampling/. Enrichment false discovery rates were controlled to 10% using the Benjamini-Hochberg correction method. Protein multifunctionality (MF) scores were also computed in ErmineJ based on the number of different GO terms with which the protein is associated. This process generates an “MF-corrected P value” (also limited to 10% FDR), which is the enrichment P value after correcting for multifunctionality.

### Differential expression of failed and healthy queens

Age-matched failing and healthy queens (11 each) were obtained from a research apiary in Pennsylvania. Queens were rated as ‘failing’ if they had ceased to lay eggs, were ‘drone layers’ (i.e., were not laying fertilized female eggs), or had otherwise inferior brood patterns. Queens were frozen on dry ice in the field and stored at −80°C until spermatheca and ovary dissection. Proteomics analysis was performed as described above. Sample handlers were blind to queen groups until all proteomics data was acquired. Differential expression analysis was performed in Perseus as described above.

## Data availability statement

All raw mass spectrometry data, protein databases, and search results are available on PRIDE ProteomeXchange (accession: PXD013728). Figures with associated raw mass spectrometry data include Figs. 4, 5, 6, and S4. Global protein abundances and p values for the laboratory heat-shock comparisons are available in Table S1. Any other data that support the findings of this study are available from the corresponding author upon request.

## Code availability statement

No specialized code central to our conclusions was used in this manuscript. R code for standard statistical analyses and figure generation will be provided upon request.

## Author correspondence

Leonard J Foster

M Marta Guarna

David R Tarpy

Jeffery S Pettis

## Acknowledgements

This work was supported by an NSERC Discovery grant (311654-11) and grants from Genome Canada and Genome British Columbia awarded to LJF, a Project Apis m grant awarded to MMG, a Project Apis m grant awarded to JSP, a USDA-NIFA grant (2016-07962) awarded to JSP and DRT, and a Boone-HodgsonWilkinson Trust grant awarded to AM and LJF. We would like to thank Alexandra Sébastien and Meghan Jelen for providing us with stink bugs and fruit flies, respectively, for the survival experiments.

We also thank Ashurst Bee Company for help with colony heat testing and Kettle Valley Queens, Nicola Valley Honey, Wild Antho, Campbells Gold Honey, Heather Meadows Honey Farm, Six Legs Good Apiaries, Wildwood Queens, Cariboo Honey, and Worker Bee Honey Company for donating failed and healthy queens for this research.

## Author contributions

AM wrote the first draft of the manuscript and revisions, conducted all data analysis, made the figures, and performed the proteomics experiments. AC and AM conducted the failed queen survey, with assistance from HH and MMG. HH and MMG executed the queen shipment temperature tracking. JM performed the survival experiments. MMG and JSP performed the drone sperm viability analyses. RU contributed the age-matched failed and healthy queens. JSP performed the queen sperm viability measurements across the range of temperatures and measured internal hive temperatures. Grants to LJF, JSP, MMG, and DRT funded the research. All authors contributed intellectually.

