## Supplemental Figures for "Honey bee queens are vulnerable to heat-induced loss of fertility"

Thermal hazards for honey bee queens

**Supplementary Material**

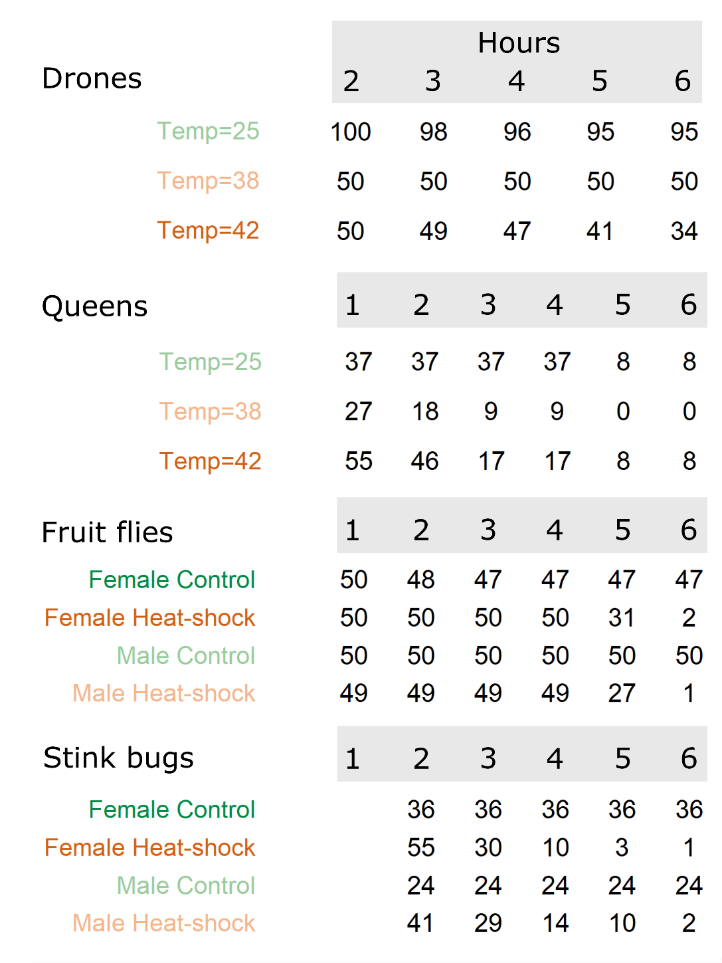

**Figure S1. Risk tables for heat-stress survival curves.** Censorship not shown. Fruit fly heat-shock was at 38 ͦC, and stink bug heat shock was at 42 ͦC.

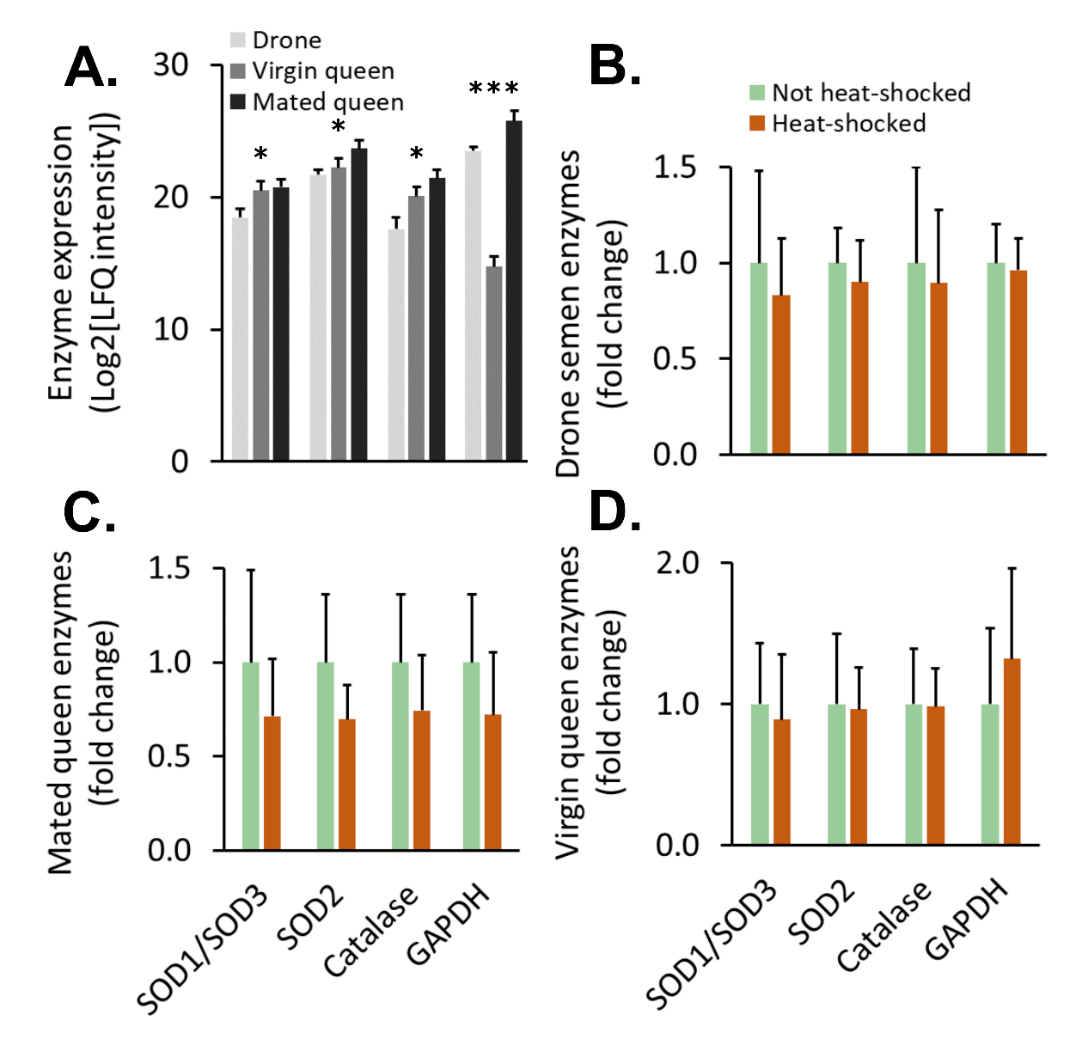

**Figure S2. Heat-shock does not affect expression of superoxide dismutase, catalase, and GAPDH.** A) Isoforms of superoxide dismutase (SOD1, SOD2, and SOD3; accessions XP_026298263.1, NP_001171519.1, and ANS15098.1, respectively), catalase (XP_026296889.1), glutathione-S-transferase (GST1-1; accession XP_026296300.1), and glyceraldehyde-3-phoshate dehydrogenase (GAPDH; accession XP_397363.4) were evaluated in drone (n = 5, where each n is composed of 4 pooled semen samples), virgin queen (n = 10), and mated queen (n = 10) reproductive tissues by intensity-based, label-free quantitative tandem mass spectrometry (permutation-based FDR = 10%). SOD1 and SOD3 were indistinguishable based on the observed peptides Error bars indicate standard deviations. * P < 1.0E-5, ***** P < 1.0E-10. B-D) Protein fold-change was evaluated between heat-shocked and non-heat-shocked treatments (n = 5 each for drones and n = 10 each for queens). Heat-shock tended to decrease enzyme abundance, but not significantly (one-way MANOVA). Error bars represent standard deviations.

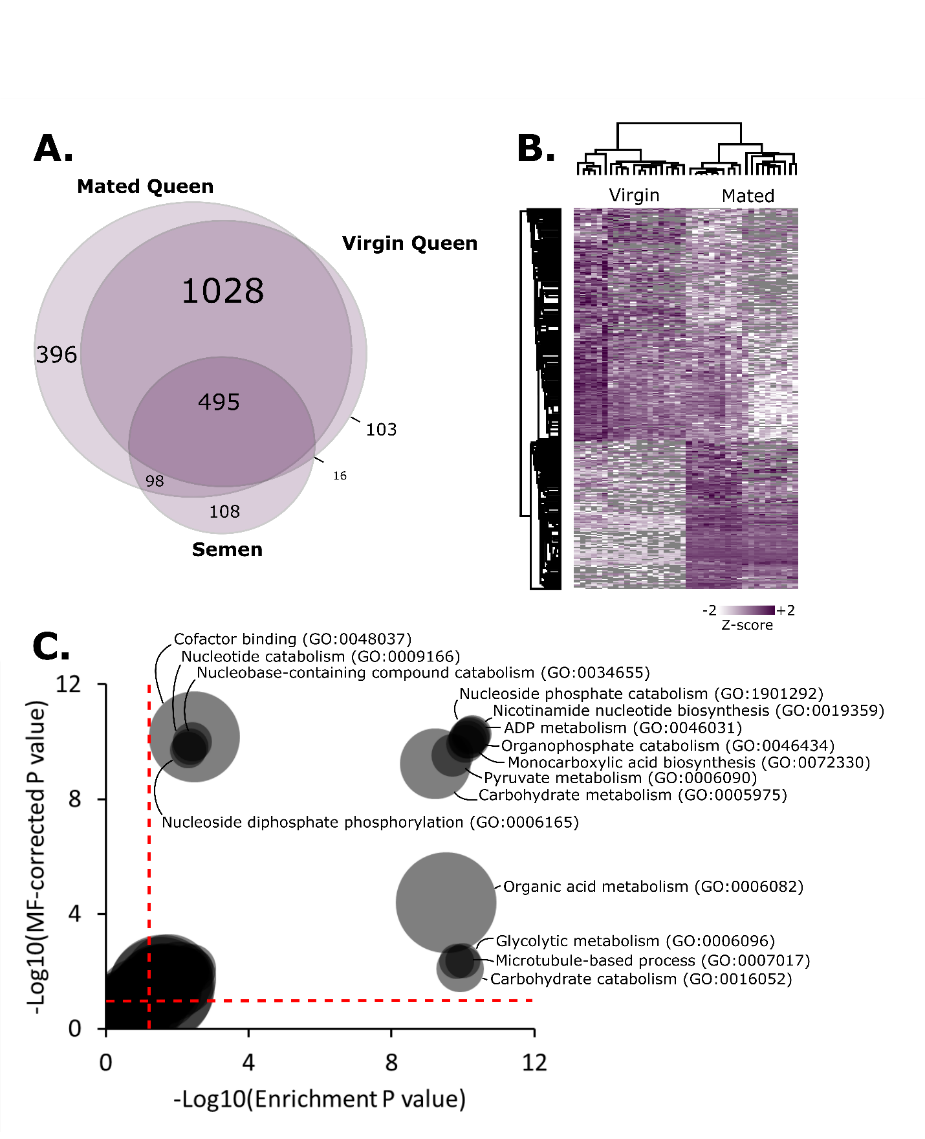

**Figure S3. Comparative proteomics of reproductive tissues.** A) Overall, 2,224 proteins were identified, including 396 proteins unique to the mated queen spermatheca, 103 unique to the virgin queen spermatheca, and 108 unique to ejaculated semen. B) 547 proteins were differentially expressed between virgin and mated queens (296 down-regulated, 250 up-regulated; 10% FDR). C) Many aspects of carbohydrate metabolism and nucleotide metabolism are significantly enriched functions (10% FDR) among the differentially expressed proteins. The x-axis depicts enrichment P values (corrected at 10% FDR) without multifunctionality correction. The y-axis depicts enrichment P values after correction for both multiple hypotheses and multifunctionality. The red dotted lines indicate the P value significance cut-off that achieves 10% FDR.

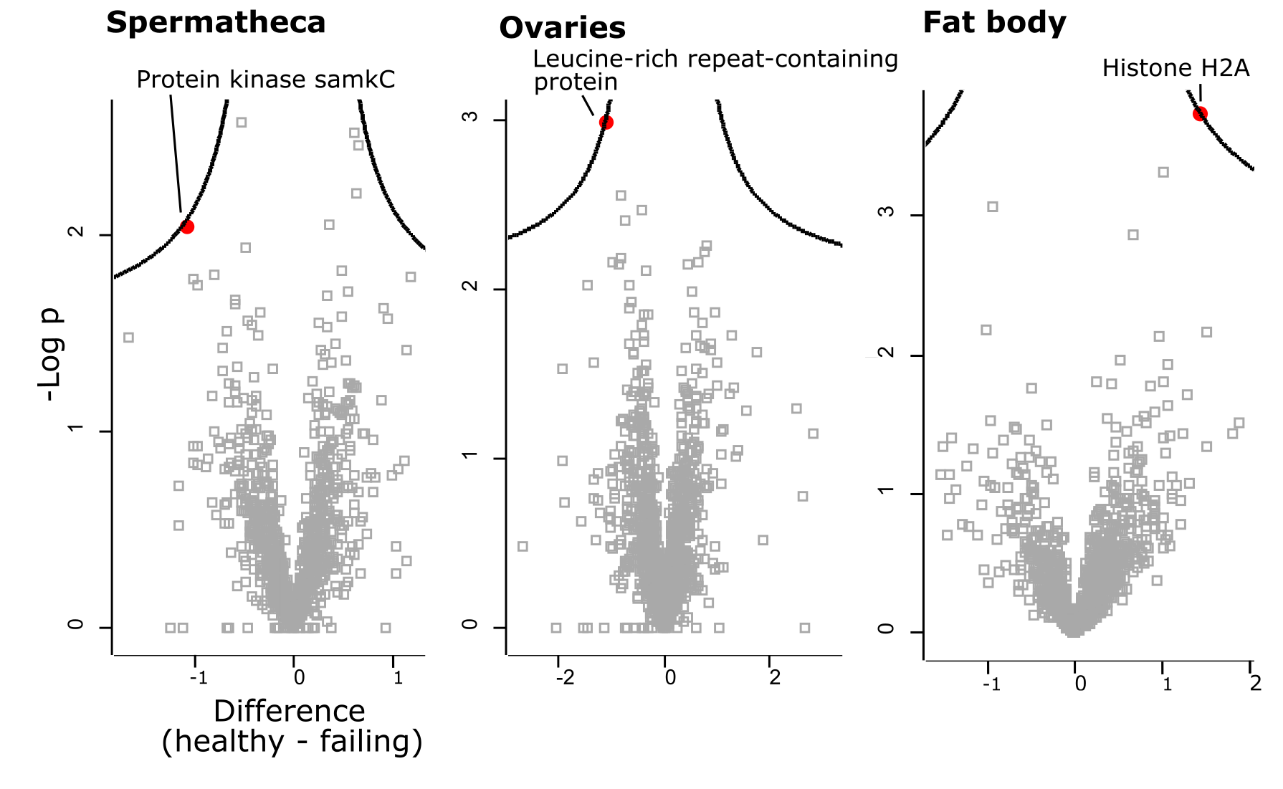

**Figure S4. Shot-gun proteomics analysis of queens heading failing and healthy colonies.** We sampled 22 queens from the field (11 heading healthy colonies, and 11 heading failing colonies) and performed quantitative proteomics analysis on their spermathecae, ovaries, and fat bodies. Only marginally statistically significant proteins were identified at 10% FDR (Benjamini-Hochberg correction).
